# The AAA-ATPase ATAD1 and its partners promote degradation of desmin intermediate filaments in muscle

**DOI:** 10.1101/2022.01.24.477493

**Authors:** Dina Aweida, Shenhav Cohen

## Abstract

Maintenance of desmin intermediate filaments (IF) is vital for muscle plasticity and function, and their perturbed integrity due to accelerated loss or aggregation causes atrophy and myopathies. Calpain-1-mediated disassembly of ubiquitinated desmin IF is a prerequisite for desmin loss, myofibril breakdown and atrophy. Because calpain-1 does not harbor a bona fide ubiquitin binding domain, the precise mechanism for desmin IF disassembly remains unknown. Here, we demonstrate that the AAA-ATPase, ATAD1, is required to facilitate disassembly and turnover of ubiquitinated desmin IF. We identified PLAA and UBXN4 as ATAD1’s interacting partners, and show they are required for desmin filaments turnover because their downregulation attenuated desmin loss upon denervation. The ATAD1-PLAA-UBXN4 complex binds desmin filaments and promotes a release of phosphorylated and ubiquitinated species into the cytosol, presenting ATAD1 as the only known AAA-ATPase that preferentially acts on phosphorylated substrates. Desmin filaments disassembly was accelerated by the coordinated functions of Atad1 and calpain-1, which interact in muscle. Thus, by extracting ubiquitinated desmin from the insoluble filament, ATAD1 may expose calpain-1 cleavage sites on desmin, consequently enhancing desmin solubilization and degradation in the cytosol.

## INTRODUCTION

Desmin intermediate filaments (IF) are critical for muscle architecture and function (Agnetti et al., 2021). By linking the contractile myofibrils to the sarcolemma and cellular organelles (i.e. mitochondria, nucleus, T-tubules, sarcoplasmic reticulum, and motor endplate (Bär et al., 2004)), this cytoskeletal network contributes to muscle structural and cellular integrity, force transmission, and mitochondrial homeostasis (Milner et al., 2000). Mutations in desmin cause myopathies in humans (Bär et al., 2004; Milner et al., 1996; Mavroidis et al., 2008) and its loss promotes muscle wasting (Helliwell et al., 1989; Boudriau et al., 1996; Aweida et al., 2018; Volodin et al., 2017; Cohen et al., 2012; Thottakara et al., 2015). Our previous studies in animal models suggested that the loss of desmin filaments in muscles atrophying due to neuronal damage (i.e. muscle denervation) (Volodin et al., 2017) or systemic catabolic states (e.g. fasting, type-2 diabetes) (Eid-Mutlak et al., 2020; Aweida et al., 2018; Cohen et al., 2012), is an initial key event triggering myofibril destruction by the ubiquitin proteasome system (UPS) (Solomon and Goldberg, 1996). The resulting excessive myofibril breakdown is a hallmark of muscle atrophy, and accounts for the associated frailty, disability and morbidity in aging or disease (Jackman and Kandarian, 2004; Cohen et al., 2014b).

We recently discovered that phosphorylation of serine residues within desmin’s head domain is required for desmin IF ubiquitination by the ubiquitin ligase TRIM32, and subsequent depolymerization (Aweida et al., 2018). Phosphorylation of desmin IF is catalyzed by glycogen synthase kinase 3-β (GSK3-β) (Kirk et al., 2014; Aweida et al., 2018), which is activated primarily by the fall in PI3K-AKT signaling (e.g. in fasting or type-2 diabetes). Although calpain-1 binds phosphorylated and ubiquitinated desmin filaments, cleavage by this enzyme probably does not require desmin ubiquitination because calpain-1 does not harbor a bona fide ubiquitin binding domain. In addition, filaments per se are not accessible to the catalytic core of the proteasome (Solomon and Goldberg, 1996), and therefore must disassemble before degradation in the cytosol (Aweida and Cohen, 2021). For example, the AAA-ATPase, p97/VCP disassembles ubiquitinated filamentous myofibrils and promotes their loss in muscles atrophying due to denervation or fasting (Piccirillo and Goldberg, 2012; Volodin et al., 2017). However, desmin IF are lost by a mechanism not requiring p97/VCP (Volodin et al., 2017). We show here that their degradation requires a distinct AAA-ATPase, ATAD1.

ATAD1 (also called THORASE) belongs to the ATPases Associated with diverse cellular Activities (AAA) superfamily of proteins that form homohexameric ring-structured complexes (Zhang and Mao, 2020; Vale, 2000). In humans, a homozygous frameshift mutation in *Atad1* causes encephalopathy, stiffness and arthrogryposis (Piard et al., 2018), but the effects of this mutation on ATAD1 function are unclear. Prior investigations have suggested a role for ATAD1 in regulation of synaptic plasticity, learning and memory (Zhang et al., 2011; Li et al., 2016), and other studies proposed that the ATAD1 yeast homolog, Msp1, is involved in the turnover of mislocalized mitochondrial membrane proteins (Li et al., 2019; Wohlever et al., 2017). Although ATAD1 appears to have important roles, its precise cellular functions remain largely obscure, and its interacting partners are not yet known. We demonstrate here that upon muscle denervation, ATAD1 catalyzes the disassembly and loss of desmin IF, which ultimately lead to myofibril breakdown and atrophy (Cohen et al., 2012; Volodin et al., 2017; Aweida et al., 2018). Phosphorylation of desmin IF is required for ATAD1 binding, presenting ATAD1 as the only known AAA-ATPase that preferentially acts on phosphorylated substrates. We identified PLAA and UBXN4 as ATAD1 interacting partners, and propose that ATAD1-PLAA-UBXN4 complex cooperates with calpain-1 to facilitate desmin IF solubilization, and subsequent degradation in the cytosol.

## RESULTS

### ATAD1 promotes muscle wasting on denervation

The Ca^2+^-dependent protease calpain-1 promotes degradation of phosphorylated and ubiquitinated desmin IF during atrophy (Aweida et al., 2018). Because calpain-1 does not harbor a bona fide ubiquitin binding domain, we sought to identify the factors that cooperate with calpain-1 in promoting desmin IF disassembly and loss. The AAA-ATPase, p97/VCP, which extracts ubiquitinated proteins from the myofibril, is not required for loss of ubiquitinated desmin filaments (Volodin et al., 2017). By incubating muscle pellets (containing insoluble desmin filaments and myofibrils) with a non-hydrolyzable ATP analogue, AMP-PNP, we recently identified enzymes that utilize ATP for their activity and act on desmin IF (Aweida et al., 2018). The addition of AMP-PNP facilitated the identification of such enzymes by mass spectrometry because they were sequestered to desmin but could not act on it (Aweida et al., 2018). The only identified AAA-ATPase enzymes on our dataset were ATAD1 and ATAD3. ATAD1 was two-fold more abundant in the insoluble fraction of 3 d denervated muscles than in innervated controls, while ATAD3 showed similar abundance in denervated and control muscles. To determine whether these AAA-ATPase enzymes also bind calpain-1, we immunoprecipitated calpain-1 from 7 d denervated muscle homogenates, when desmin depolymerization by calpain-1 is accelerated (Volodin et al., 2017; Aweida et al., 2018), and analyzed protein precipitates by mass spectrometry. Interestingly, ATAD1 was the only AAA-ATPase that bound calpain-1, and was also one of the most abundant proteins in the sample (1000 times more abundant in the calpain-1 immunoprecipitation sample than in IgG control). These findings were unexpected because ATAD1 has been considerably understudied, and there had been no prior reports of ATAD1 in muscle. In denervated skeletal muscle, however, ATAD1 binds both the insoluble filaments and calpain-1, and is critical for desmin filament loss and atrophy (see below).

To clarify the role of ATAD1, we first determined whether it is induced during atrophy. The levels of mRNA for *Atad1* increased significantly at 7 and 10 d after denervation (Fig. 1B), just when desmin depolymerization by calpain-1 is rapid (Aweida et al., 2018), but returned to basal levels at 14 d, when the content of phosphorylated and ubiquitinated desmin filaments is substantially low and myofibril disassembly by p97/VCP is accelerated (Volodin et al., 2017). To test whether ATAD1 is important for atrophy, we suppressed its expression by electroporation of shRNA plasmid into mouse Tibialis Anterior (TA) muscle. Two shRNAs had been generated, shAtad1-1 and shAtad1-2, which reduced *Atad1* mRNA levels in mouse muscles below the levels in shLacz expressing controls (Fig. 1B), and here we used the potent shAtad1-1. The downregulation of *Atad1* was sufficient to attenuate muscle fiber atrophy because at 14 d after denervation the cross-sectional area of 975 fibers expressing shAtad1 was significantly larger than that of 975 non-transfected denervated fibers in the same muscle (n=8 mice) (Fig. 1C and Table I). Therefore, ATAD1 is important for muscle atrophy induced by denervation.

**Figure 1:**
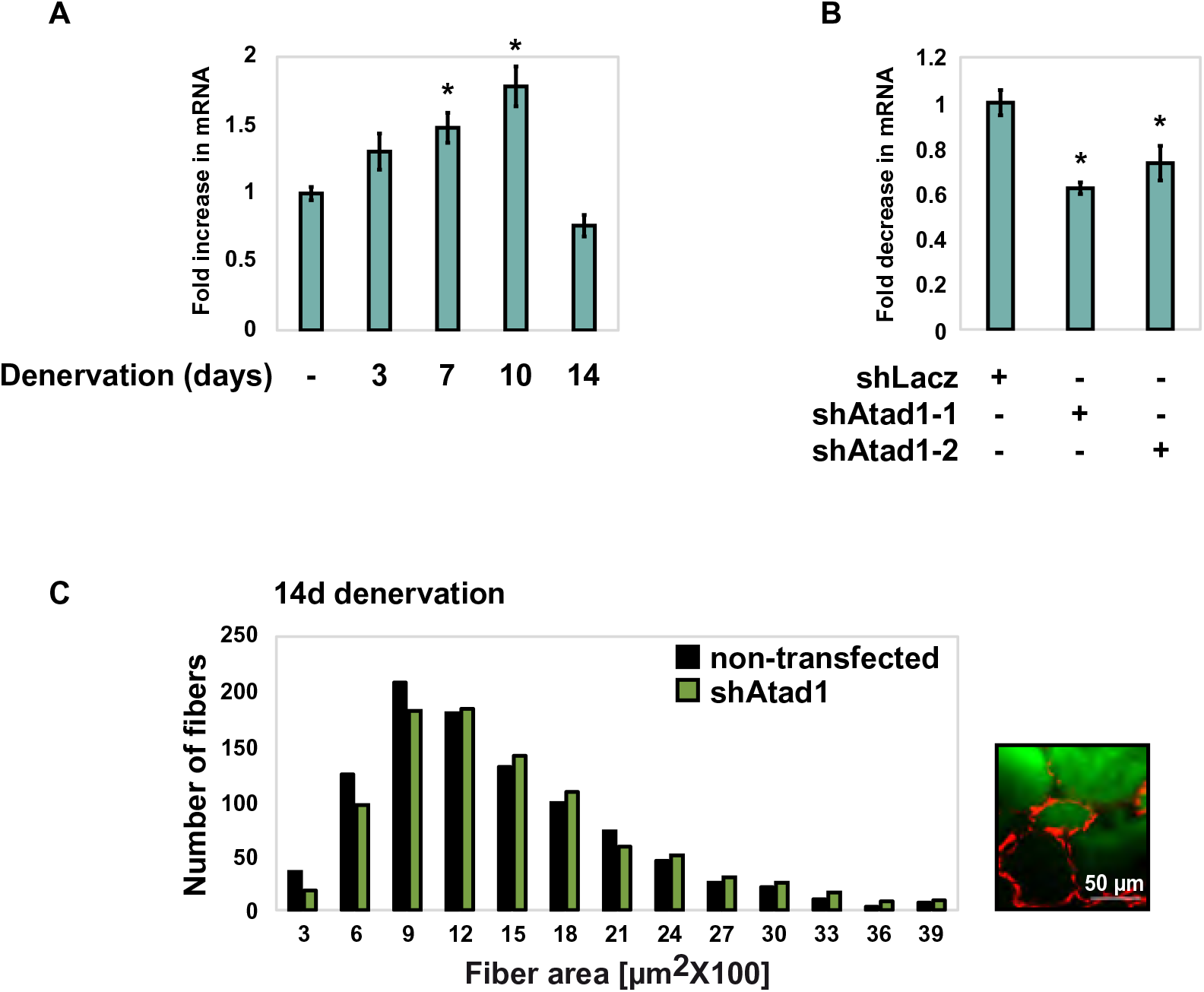
ATAD1 promotes muscle atrophy on denervation. (A) *Atad1* is induced upon muscle denervation. RT-PCR of mRNA preparations from innervated muscles and ones denervated for 3, 7, 10, and 14 d using specific primers for *Atad1*. Data are plotted as the mean fold change relative to innervated control ± SEM. n = 4. *, P < 0.05 *vs.* innervated. (B) shRNA-mediated knockdown of *Atad1* in 14 d denervated TA muscles. RT-PCR of mRNA preparations from 14 d denervated muscles expressing shLacz control or two different shRNAs against *Atad1*, shAtad1-1 and shAtad1-2, using specific primers for *Atad1*. Data are plotted as the mean fold change relative to shLacz control ± SEM. n = 4. *, P < 0.05 *vs.* shLacz. (C) *Atad1* downregulation attenuates fiber atrophy on denervation. Measurements of cross-sectional areas of 973 fibers expressing shAtad1 (green bars) versus 973 non-transfected fibers (black bars) in the same muscle. n = 8 mice. Bar: 50 µm.

**Table I.**
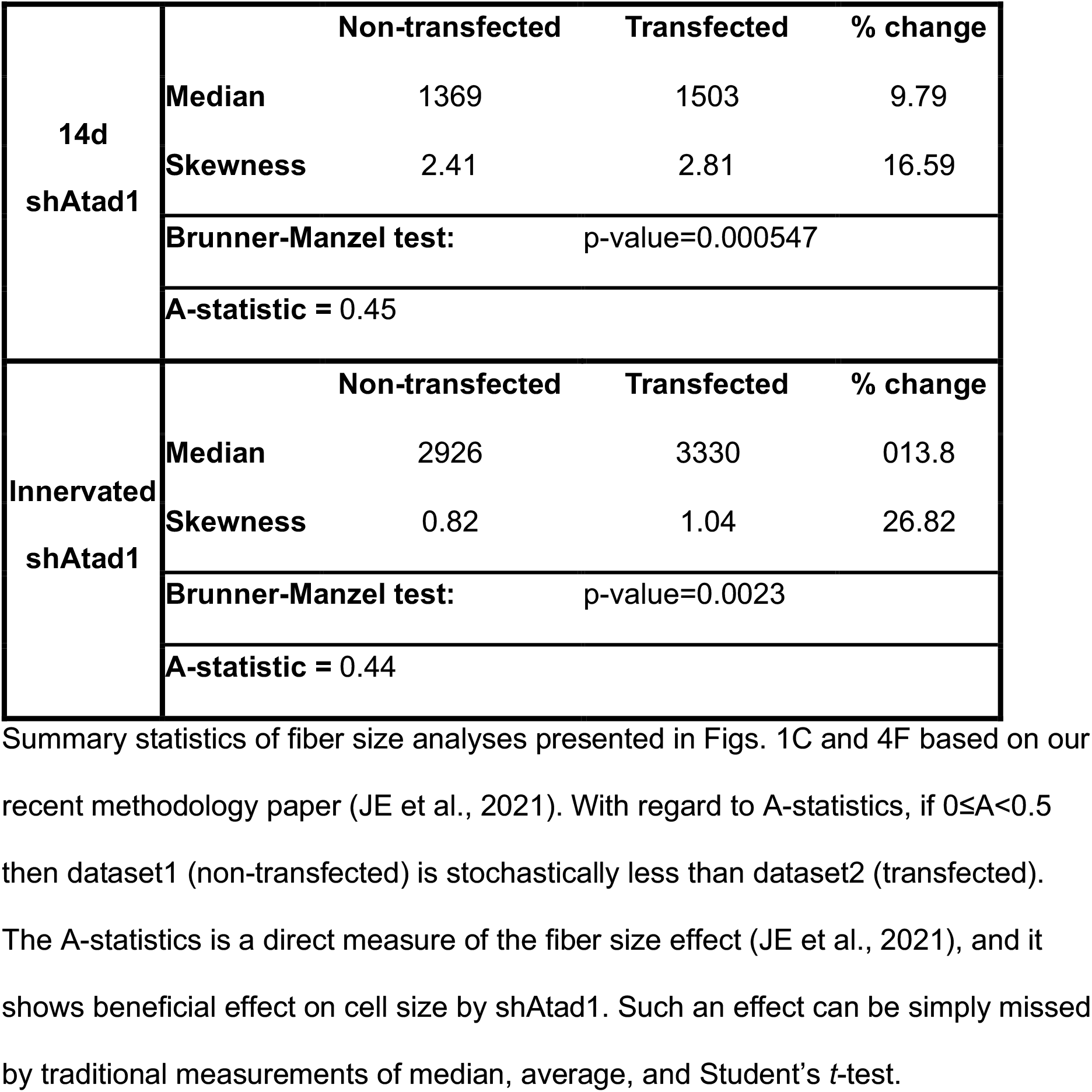

### ATAD1 binds phosphorylated desmin filaments and promotes their disassembly

To investigate if this reduction in fiber atrophy occurred through the attenuation of desmin IF loss, which should reduce overall proteolysis (Cohen, 2020; Aweida and Cohen, 2021; Agnetti et al., 2021), we initially confirmed our mass spectrometry data (Aweida et al., 2018) and determined whether ATAD1 in fact binds desmin filaments *in vivo*. For this purpose, we isolated desmin filaments from muscles at different times after denervation, and analyzed by SDS-PAGE and immunoblotting. Ubiquitination of desmin IF increased at 3 d after denervation (Fig. 2A), as we had previously shown (Volodin et al., 2017), but was markedly reduced at 14 d after denervation, when ubiquitinated desmin is degraded (Volodin et al., 2017). It is noteworthy that the amount of ubiquitinated desmin IF at 7 d after denervation did not differ significantly from that in innervated controls (Fig. 2A) because at this time after denervation depolymerization of ubiquitinated desmin IF is accelerated (Volodin et al., 2017). Interestingly, at 3 d after denervation, when desmin filaments are rapidly ubiquitinated, ATAD1 accumulated on desmin IF (Fig. 2B). However, at 7 d after denervation, when desmin IF depolymerization by calpain-1 is accelerated (Aweida et al., 2018; Volodin et al., 2017), ATAD1 association with desmin filament was markedly reduced; instead, ATAD1 mainly accumulated in the cytosol (Fig. 2B), most likely bound to ubiquitinated species of desmin (see below).

**Figure 2:**
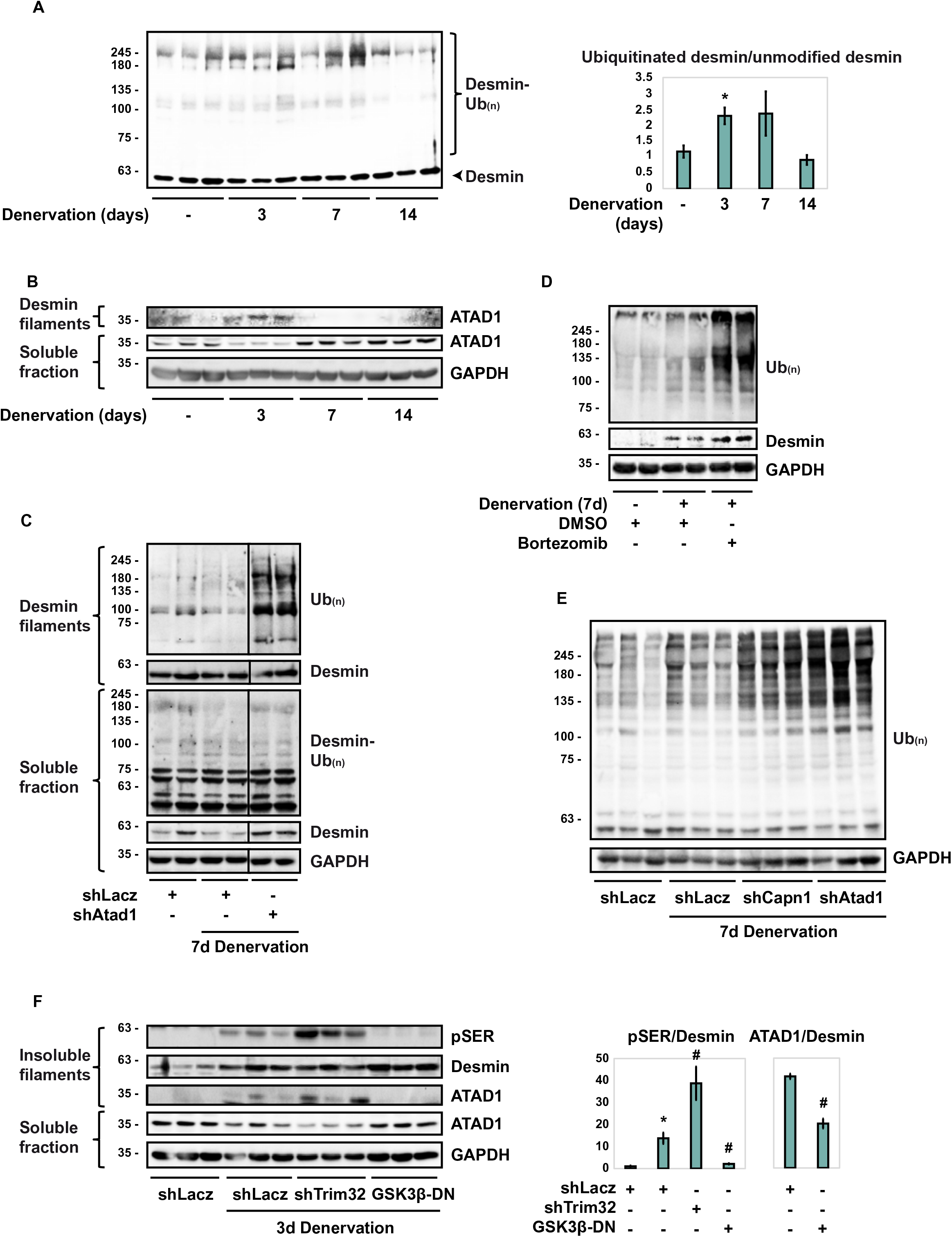
ATAD1 binds phosphorylated desmin filaments and promotes their disassembly. (A) Desmin ubiquitination increases at 3 d after denervation. Left: desmin filaments isolated from muscles at 0, 3, 7, or 14 d after denervation were analyzed by immunoblotting. Right: densitometric measurement of presented blots. Mean ratios of ubiquitinated desmin to unmodified desmin ± SEM are depicted in a graph. n = 3 mice. *, P < 0.05 *vs.* innervated. (B) ATAD1 binds desmin filaments at 3 d after denervation, and at 7 d after denervation it is released from the pellet and accumulates in the soluble fraction. Desmin filaments and soluble fractions isolated from innervated and denervated muscles were analyzed by immunoblotting. (C) Downregulation of *Atad1* prevents depolymerization of ubiquitinated desmin filaments. Desmin filaments and soluble fractions from innervated and 7 d denervated TA muscles expressing shLacz or shAtad1 were analyzed by immunoblotting. Black line indicates the removal of intervening lanes for presentation purposes. (D) Desmin is degraded by the proteasome in atrophying muscles. Soluble fractions from innervated and denervated (7 d) muscles of mice injected with Bortezomib or DMSO were analyzed by SDS-PAGE and immunoblotting. (E) *Atad1* downregulation results in accumulation of ubiquitinated soluble proteins. Soluble fractions from innervated and denervated muscles expressing shLacz, shCapn1 or shAtad1 were analyzed by SDS-PAGE and immunoblotting. (F) Desmin filament phosphorylation is a prerequisite for ATAD1 binding. Left: insoluble and soluble fractions from innervated or denervated muscles expressing shLacz, shTrim32 or GSK3-β-DN were analyzed by immunoblotting. ATAD1 was detected on isolated desmin filaments. Right: densitometric measurement of presented blots. Mean ratios of pSER to desmin ± SEM are depicted in a graph. n = 3. *, P < 0.05 *vs*. shLacz in innervated; #, P < 0.05 *vs.* shLacz in denervated.

To test more directly whether ATAD1 is required for desmin filament depolymerization and loss, we employed a similar approach as we used to analyze disassembly of filamentous myofibrils by p97/VCP (Volodin et al., 2017). We determined the effects of ATAD1 downregulation on the total amount of ubiquitinated desmin in the soluble and insoluble fractions of 7 d denervated muscles, just when disassembly of desmin IF is accelerated (Volodin et al., 2017). Downregulation of ATAD1 with shAtad1 prevented disassembly of desmin IF because ubiquitinated desmin accumulated as insoluble filaments (Fig. 2C). In fact, the amount of ubiquitinated filaments in these shAtad1-expressing muscles exceeded the amounts observed in shLacz expressing denervated controls. Thus, the ubiquitination of desmin IF increases on denervation (Fig. 2A and (Volodin et al., 2017)), and ATAD1 catalyzes the solubilization and degradation of these ubiquitinated proteins.

In addition, desmin was released from the cytoskeleton into the cytosol as ubiquitinated species, and it was then degraded (Fig. 2C), but not when endogenous ATAD1 was downregulated with shAtad1 (Fig. 2C). Degradation of soluble desmin was validated *in vivo* by the injection of mice with the proteasome inhibitor, Bortezomib (3mg/kg body weight), which prevented desmin degradation in denervated (7 d) muscles and led to accumulation of desmin in its ubiquitinated form mediated (compare to DMSO injected controls, Fig. 2D). Interestingly, downregulation of either *Atad1* or calpain1 (shCapn1 is described in (Aweida et al., 2018)) led to a dramatic increase in the content of overall ubiquitin conjugates in the cytosol (Fig. 2E), indicating that on denervation, degradation of soluble proteins also depends on ATAD1 and calpain1 functions. Thus, ubiquitinated desmin seems to be released from the filament into the cytosol for degradation, and this step requires ATAD1.

ATAD1 binds desmin IF early after denervation (3 d), when desmin filaments are phosphorylated by GSK3-β and then ubiquitinated by the ubiquitin ligase, TRIM32 (Aweida et al., 2018). To determine if these post-translational modifications of desmin filaments are required for association with ATAD1, we purified desmin filaments from innervated and 3 d denervated muscles expressing GSK3-β dominant negative (GSK3-β-DN), *Trim32* shRNA (shTrim32), or shLacz control and analyzed by immunoblotting (GSK3-β-DN and shTrim32 were described in (Aweida et al., 2018; Volodin et al., 2017; Cohen et al., 2014a)). At 3 d after denervation desmin phosphorylation increased and inhibition of GSK3-β with GSK3-β-DN prevented this phosphorylation (Fig. 2F), as reported (Aweida et al., 2018). Loss of desmin IF by TRIM32-mediated ubiquitination requires desmin phosphorylation (Aweida et al., 2018; Volodin et al., 2017; Cohen et al., 2012). Consistently, downregulation of *Trim32* prevented this loss, and desmin accumulated as phosphorylated filaments (Fig. 2F). Desmin IF phosphorylation seems to be important for ATAD1 binding because the association of ATAD1 with desmin increased correlatively as phosphorylated desmin filaments accumulated, if by the induction of atrophy or by the downregulation of *Trim32* (Fig. 2F). Moreover, ATAD1 was less bound to desmin in muscles where GSK3-β was inactive (Fig. 2F). Therefore, desmin IF phosphorylation is a prerequisite for ATAD1 binding.

### Isolation of ATAD1 binding partners from muscle homogenates

To investigate if ATAD1 cooperates with adaptor proteins to promote desmin IF disassembly, we searched for its interacting partners *in vivo* using three independent mass spectrometry-based proteomic approaches. The first approach we employed was size-exclusion chromatography (SEC) of 7 d denervated muscles because at this time after denervation ubiquitinated desmin IF are solubilized by calpain-1 (Volodin et al., 2017; Aweida et al., 2018), and ATAD1 accumulates in the soluble phase (Fig. 2B) bound to calpain-1 (see also below Fig. 3D). To facilitate the identification of low abundant proteins, mouse muscles (1 g) were cross-linked in 1% PFA, homogenized, and after 10 min of incubation the cross-linking reaction was quenched with glycine (0.125M). Ammonium sulfate precipitates were then solubilized and fractionated by a gel filtration column (Superdex 200 10/300 GL), and protein eluates were analyzed by SDS-PAGE, immunoblotting, and silver staining (Fig. 3A-B). ATAD1 was predominantly recovered in fraction #8, together with desmin and additional low and high molecular weight proteins (Fig. 3B). Mass spectrometry analysis of this fraction in two biological replicates of denervated muscles identified ATAD1 and several UPS components, including E1 and ubiquitin, E2s, both RING and HECT domain E3s, proteasome subunits, deubiquitinating enzymes, calpain 1 and 2, and SUMO activating enzyme (Table II). Only UPS components that were identified with ≥ 2 unique peptides were categorized based on function using DAVID annotation tool.

**Figure 3:**
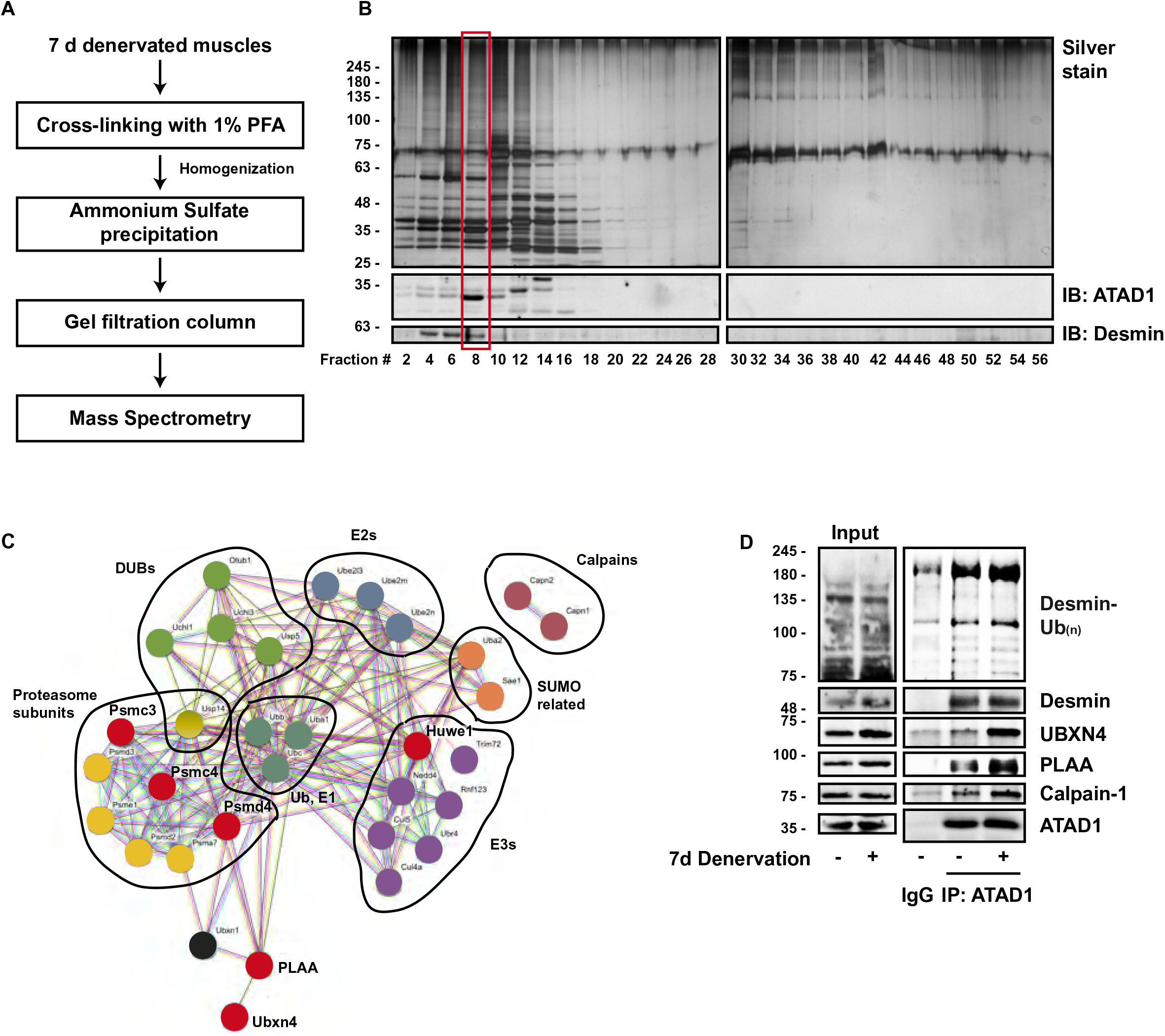
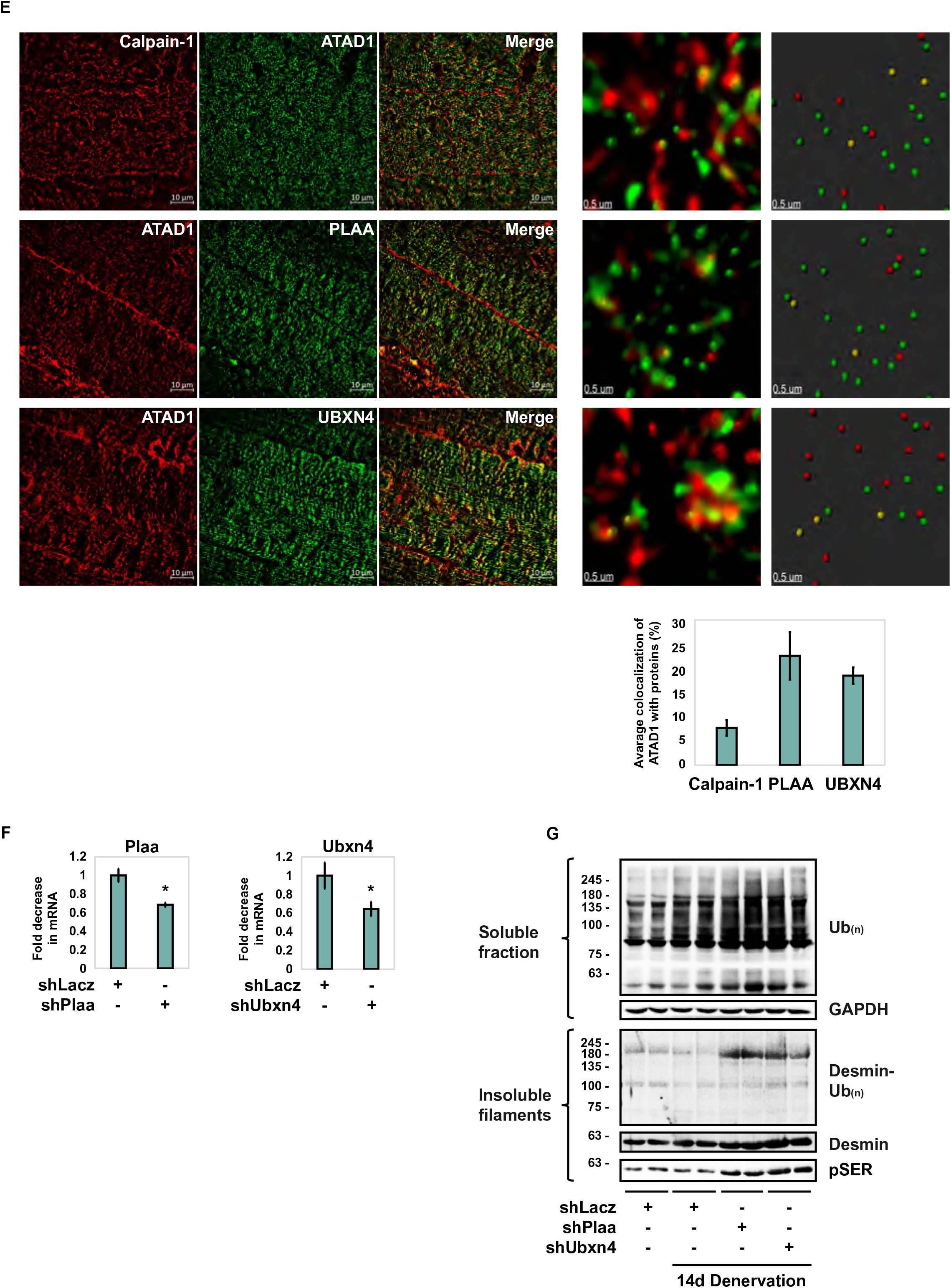
ATAD1-PLAA-UBXN4 complex promotes disassembly of ubiquitinated desmin filaments. (A) Scheme of ATAD1’s purification from muscle and isolation of its binding partners. (B) Analysis of size-exclusion chromatography fractions by SDS-PAGE, silver staining (top panel) or immunoblotting (lower panel). (C) Interaction networks for UPS components and related proteins identified by three independent mass spectrometry-based proteomics approaches using the STRING database. UPS enzymes are grouped based on function. Labeled in red are proteins that were at least 3 times more abundant on desmin filaments at 3 d after denervation. (D) ATAD1 and its partners bind ubiquitinated desmin, and these associations increase after denervation. ATAD1 was immunoprecipitated from innervated and 7 d denervated muscle homogenates, and protein precipitates were analyzed by immunoblotting. (E) ATAD1 colocalizes with calpain-1, PLAA and UBXN4 in muscle. Left: Structured illumination microscopy (SIM) images (bar, 10 μm) of 7 d denervated muscle longitudinal sections stained with the indicated antibodies. Right: representative analysis of SIM images using the spots module of the Imaris software (bar, 0.5 μm). Only spots that were within a distance threshold of less than 100 nm were considered colocalized. Bottom: average colocalization of ATAD1 with the indicated proteins. Data are represented as mean ± SEM. n=3. (F) shRNA-mediated knockdown of *Plaa* and *Ubxn4* in HEK293 cells. RT-PCR of mRNA preparations from HEK293 cells expressing shLacz, shPlaa, or shUbxn4 using specific primers for *PLAA* or *UBXN4*. Data are plotted as the mean fold change relative to control ± SEM. n = 5. *, P < 0.05 *vs.* shLacz. (G)Downregulation of *Plaa* or *Ubxn4* prevents the loss of desmin and soluble proteins in denervated muscle. Insoluble and soluble fractions from atrophying muscles expressing shLacz, shPlaa or shUbxn4 were analyzed by immunoblotting.

**Table II.** ATAD1 binds a variety of UPS components in muscle. ATAD1 interacting partners were identified by three independent mass spectrometry-based proteomics approaches. Only UPS components that were identified with ≥ 2 unique peptides were categorized based on function using DAVID annotation tool. Two proteomics approaches were oriented specifically towards identifying ATAD1-binding partners: SEC (427 total proteins were identified), and Atad1 immunoprecipitation (592 total proteins were identified). These lists of UPS components were compared to our previous kinase-trap assay dataset ((Aweida et al., 2018), 1552 total proteins were identified) and the number of unique peptides only for proteins that overlapped is listed here. The kinase trap assay was used to identify proteins that utilize ATP for their function and act on desmin, and ATAD1 was one of the most abundant proteins in the sample.

|  |  | #Unique peptides |  |  |  |
| --- | --- | --- | --- | --- | --- |
| Protein | Gene | SEC | ATAD<br>1 IP | Bound to<br>desmin<br>filaments at<br>3d after<br>denervation | Fold change<br>(3d<br>denervated/<br>innervated) |
| Ubiquitin-like<br>modifier-activating<br>enzyme 1 | *UBA1* | 37 |  | 3 |  |
| Ubiquitin-<br>conjugating enzyme<br>E2 L3 | *UBE2L3* | 3 | 2 | 5 |  |
| Ubiquitin-<br>conjugating enzyme<br>E2 N | *UBE2N* | 2 |  | 5 |  |
| NEDD8-conjugating<br>enzyme Ubc12 | *UBE2M* | 2 |  | 7 |  |
| E3 ubiquitin-protein<br>ligase UBR4 | *UBR4* |  | 4 | 86 |  |
| E3 ubiquitin-protein<br>ligase NEDD4 | *NEDD4* | 4 | 5 |  |  |
| Tripartite motif-<br>containing protein<br>72 | *TRIM72* | 2 | 10 | 40 |  |
| E3 ubiquitin-protein<br>ligase RNF123 | *RNF123* | 19 | 3 | 6 |  |
| E3 ubiquitin-protein<br>ligase HUWE1 | *HUWE1* |  | 5 | 50 | 3.7 |
| Cullin-5 | *CUL5* |  | 6 | 6 |  |
| Cullin-4A | *CUL4A* |  | 2 |  |  |
| Polyubiquitin-B | *UBB* | 6 |  | 7 |  |
| Polyubiquitin-C | *UBC* |  | 5 | 6 |  |
| 26S proteasome<br>non-ATPase<br>regulatory subunit 4 | *PSMD4* | 4 |  | 5 | 4.5 |
| Proteasome subunit<br>alpha type-7 | *PSMA7* | 3 |  | 5 |  |
| Proteasome<br>activator complex<br>subunit 1 | *PSME1* | 3 |  |  |  |
| 26S protease<br>regulatory subunit<br>6A | *PSMC3* |  | 2 | 24 | 8.6 |
| 26S proteasome<br>regulatory subunit<br>6B | *PSMC4* |  | 4 | 24 | 6.7 |
| 26S proteasome<br>non-ATPase<br>regulatory subunit 2 | *PSMD2* |  | 4 | 3 |  |
| 26S proteasome<br>non-ATPase<br>regulatory subunit 3 | *PSMD3* |  | 4 | 37 |  |
| Ubiquitin carboxyl-<br>terminal hydrolase<br>14 | *USP14* | 4 |  | 4 |  |
| Ubiquitin carboxyl-<br>terminal hydrolase 5 | *USP5* | 3 |  | 2 |  |
| Ubiquitin carboxyl-<br>terminal hydrolase<br>isozyme L1 | *UCHL1* | 2 |  |  |  |
| Ubiquitin carboxyl-<br>terminal hydrolase<br>isozyme L3 | *UCHL3* | 2 |  |  |  |
| Ubiquitin<br>thioesterase OTUB1 | *OTUB1* |  | 2 | 7 |  |
| Calpain-1 catalytic<br>subunit | *CAPN1* | 12 | 12 | 24 |  |
| Calpain-2 catalytic<br>subunit | *CAPN2* | 2 | 2 | 18 |  |
| SUMO-activating<br>enzyme subunit 1 | *SAE1* | 2 |  |  |  |
| SUMO-activating<br>enzyme subunit 2 | *UBA2* | 3 |  |  |  |
| Phospholipase A-2-activating protein | *PLAA* |  | 4 | 13 | 4.9 |
| UBX domain-containing protein 4 | *UBXN4* |  |  | 4 | 4.7 |
| UBX domain-containing protein 1 | *UBXN1* | 2 |  |  |  |

To confirm these findings by an independent approach, we immunoprecipitated ATAD1 from 3 d denervated muscle homogenates (6000 x *g* supernatant), and washed the resulting precipitates extensively with buffer containing 500 mM NaCl to remove nonspecific or weakly associated proteins. The ATAD1-bound proteins were identified by mass spectrometry, which revealed several of the aforementioned components, and additional UPS components as well as Phospholipase A-2-activating protein (PLAA) (Doa1/Ufd3 in yeast) (Table II), which contains WD40 repeats that can bind ubiquitinated proteins (Pashkova et al., 2010).

To corroborate these findings further, we employed an additional strategy and determined if any of these proteins accumulate on desmin filaments together with ATAD1 at 3 d after denervation (Fig. 2B). As mentioned above, using AMP-PNP and mass spectrometry, we previously identified enzymes that utilize ATP for their activity and bind with high affinity to desmin filaments (Aweida et al., 2018). Searching this list of enzymes, we found an overlap with almost all proteins identified by SEC and Atad1 immunoprecipitation analyses (Fig. 3A-C and Table II). Using the STRING database, we generated a network based on established annotation for protein interactors and found a significant interconnectivity among the vast majority of the identified enzymes. The exception was calpain-1 and −2, which have no known interactions with the other components on our dataset (Fig. 3C). Here we present evidence that calpain-1 interacts with several components on this network to promote disassembly of desmin filaments (see below Fig. 3D).

Out of the 32 identified proteins in our dataset, six were at least threefold more abundant on desmin filaments at 3d after denervation (Table II, and labeled in red in Fig. 3C), the time when ATAD1 accumulates on desmin (Fig. 2B), including 3 proteasome subunits PSMD4 (Rpn10), PSMC4 (Rpt3), and PSMC3 (Rpt5), the ubiquitin ligase HUWE1, PLAA, and UBXN4 (UBXD2/erasin), which contains a Ubiquitin regulatory X (UBX) domain and is known to promote recruitment of ubiquitinated substrates to AAA-ATPase complexes (e.g., p97/VCP (Lim et al., 2009)).

### Downregulation of *Plaa* or *Ubxn4* prevents desmin solubilization and degradation, and loss of soluble ubiquitinated proteins

To clarify the mechanism for ATAD1-mediated disassembly of phosphorylated and ubiquitinated desmin filaments, we focused our attention on UBXN4 and PLAA because these components mediate degradation of ubiquitinated proteins (Lim et al., 2009; Ren et al., 2008), and harbor ubiquitin binding domains (WD40 repeats in PLAA and UBX domain in UBXN4). We initially validated our mass spectrometry data and investigated if UBXN4 and PLAA in fact form an intact complex with ATAD1 in muscle. UBXN4 and PLAA could be coprecipitated with ATAD1 from innervated muscle, and these associations increased after denervation (Fig. 3D). This protein assembly, which also contained calpain-1, bound ubiquitinated desmin filaments, and muscle denervation enhanced these associations (Fig. 3D), suggesting that UBXN4 and PLAA may be adaptor proteins that link ubiquitinated desmin to ATAD1 complex to promote desmin IF disassembly. These new associations were further validated by an immunofluorescence staining of longitudinal sections from 7 d denervated muscles and super-resolution Structured illumination microscopy (SIM), which demonstrated colocalization of ATAD1 with calpain-1, PLAA and UBXN4 (Fig. 3E). To confirm that these proteins in fact colocalize, we measured the average colocalization of ATAD1 with calpain-1, PLAA and UBXN4 using the spots detection and colocalization analysis of the Imaris software (Fig. 3E). Only spots that were within a distance threshold of less than 100 nm were considered colocalized (Fig. 3E, graph).

To determine if PLAA and UBXN4 are required for desmin IF loss, we analyzed the reduction in the amount of ubiquitinated desmin filaments upon denervation after electroporation of PLAA and UBXN4 shRNAs (shPlaa and shUbxn4, respectively), which efficiently reduced *Plaa* and *Ubxn4* expression (Fig. 3F). In accord with prior findings (Volodin et al., 2017), the amount of ubiquitinated desmin filaments decreased in muscle at 14 d after denervation due to their degradation (Fig. 3G). However, downregulation of either *Plaa* or *Ubxn4* blocked this decrease, and instead desmin accumulated as insoluble phosphorylated and ubiquitinated filaments, which exceeded the amounts observed in innervated muscles (Fig. 3G). Thus, phosphorylation and ubiquitination of desmin IF increase on denervation, as reported (Volodin et al., 2017), and PLAA and UBXN4 promote the solubilization and degradation of these filaments.

Our prior findings imply that stabilization of desmin filaments attenuates overall proteolysis (Aweida et al., 2018; Volodin et al., 2017). Accordingly, the muscle’s content of soluble ubiquitin conjugates increased at 14 d after denervation compared with innervated controls, as we had shown (Volodin et al., 2017). However, the downregulation of *Plaa* or *Ubxn4* caused a more dramatic increase in the total content of ubiquitin conjugates to levels that exceeded those in denervated muscles (Fig. 3G), indicating that, similar to ATAD1 (Fig. 2E), PLAA and UBXN4 are also required for degradation of proteins in the cytosol.

### ATAD1 and calpain-1 cooperatively promote desmin filament disassembly and loss

Because ATAD1 in denervated muscles was bound to calpain-1, and since both enzymes promote dissociation of phosphorylated desmin filaments, we investigated whether they cooperate in promoting desmin IF loss. Muscles were electroporated with either shCapn1 or shAtad1, or with both constructs (Fig. 4A-B), and the insoluble fraction was analyzed by SDS-PAGE and immunoblotting (Fig. 4C). Expressing either shCapn1 or shAtad1 attenuated disassembly and loss of phosphorylated desmin IF on denervation, and desmin accumulated as insoluble phosphorylated filaments (Fig. 4C). Interestingly, this beneficial effect on desmin was much larger by simultaneously downregulating both enzymes, which caused a more dramatic accumulation of phosphorylated desmin IF (Fig. 4C). Simultaneous downregulation of both *Atad1* and calpain-1 also affected soluble ubiquitin conjugates, which accumulated in the cytosol to a greater extent than when each enzyme was downregulated alone (Fig. 4D). Thus, ATAD1 and calpain-1 cooperate in an additive fashion in promoting desmin IF loss and protein degradation during atrophy.

**Figure 4:**
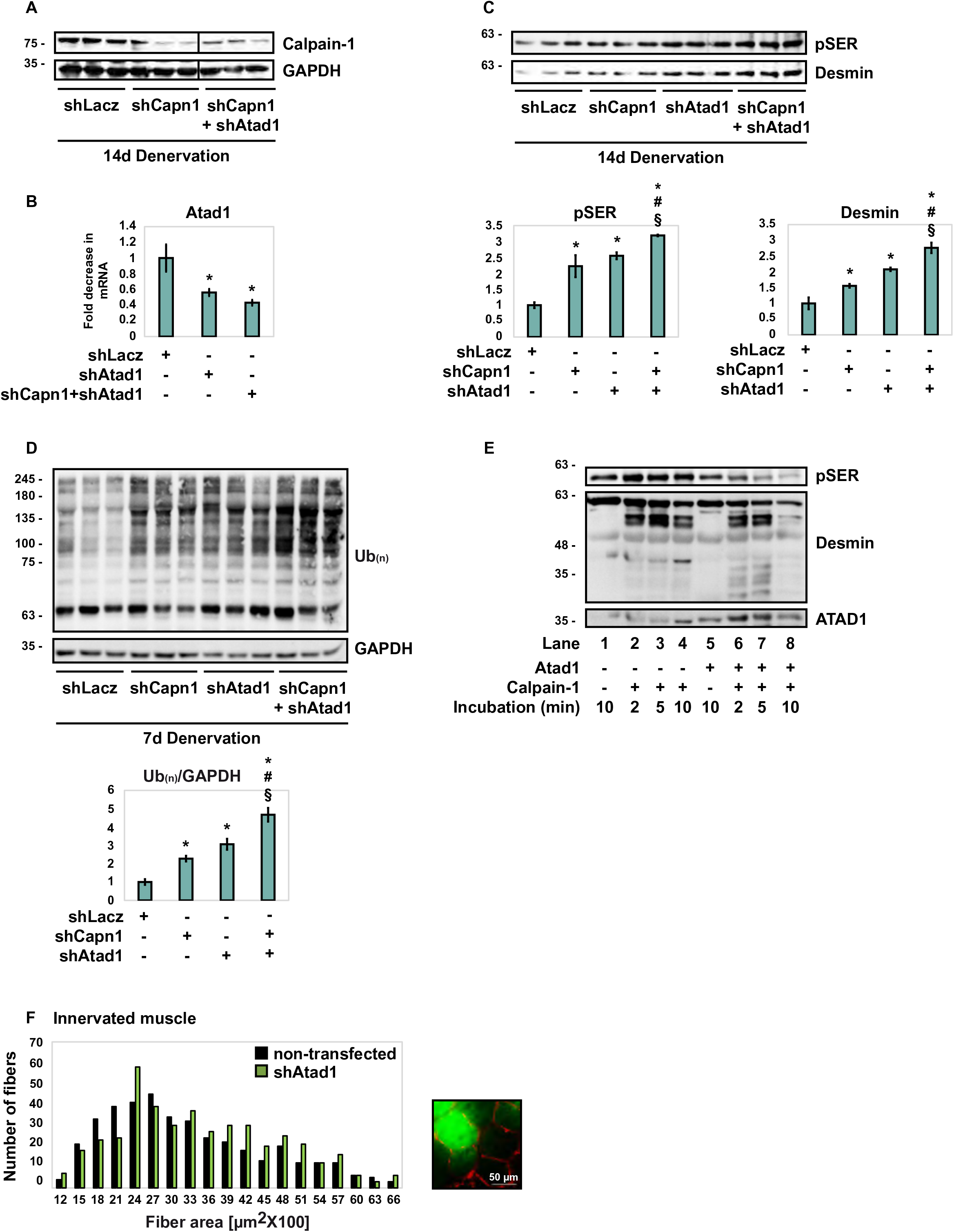
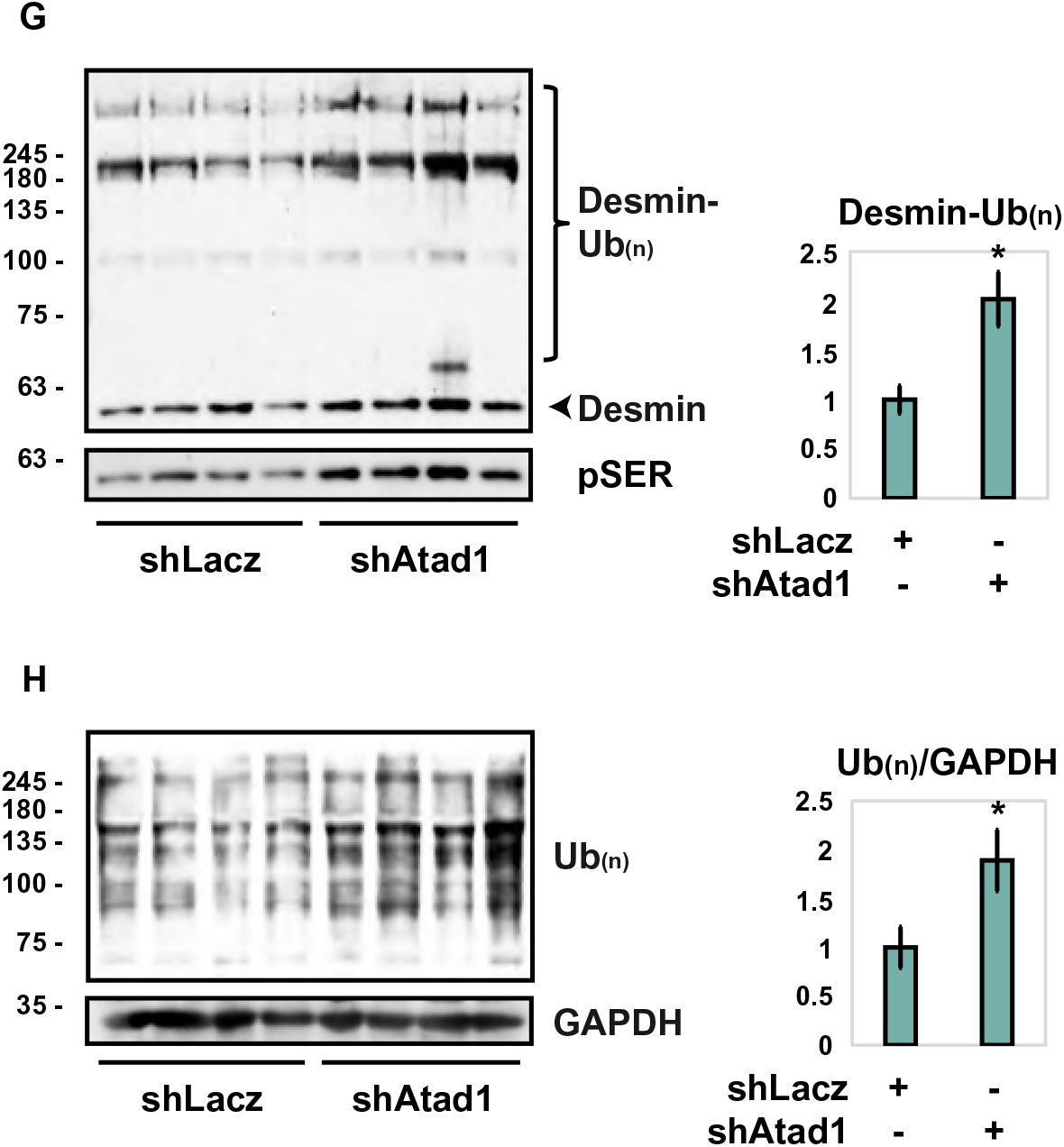
ATAD1 and calpain-1 cooperatively promote desmin filament disassembly and loss. (A) shRNA-mediated knockdown of calpain-1 in 14 d denervated TA muscles. Homogenates from denervated muscles expressing shLacz, shCapn1 or co-expressing shCapn1 and shAtad1 were analyzed by immunoblotting. Black line indicates the removal of intervening lanes for presentation purposes. (B) shRNA-mediated knockdown of *Atad1* in 14 d denervated TA muscles. RT-PCR of mRNA preparations from denervated muscles expressing shLacz, shCapn1 or co-expressing shCapn1 and shAtad1 using specific primers for *Atad1*. Data are plotted as the mean fold change relative to control ± SEM. n = 4. *, P < 0.05 *vs.* denervated shLacz. (C) On denervation, ATAD1 and calpain-1 cooperate in an additive fashion in promoting desmin IF. Top: desmin filaments isolated from denervated muscles expressing shLacz, shCapn1, shAtad1 or co-expressing shCapn1 and shAtad1 were analyzed by immunoblotting. Bottom: densitometric measurement of presented blots. Mean ± SEM. n = 3. *, P < 0.05 *vs.* shLacz; #, P < 0.05 *vs.* shCapn1 in atrophy; §, P < 0.05 *vs.* shAtad1 in atrophy. (D) ATAD1 and calpain-1 cooperate in an additive fashion in promoting loss of soluble proteins in denervated muscle. Left: homogenates from denervated muscles expressing shLacz, shCapn1, shAtad1 or co-expressing shCapn1 and shAtad1 were analyzed by immunoblotting. Bottom: densitometric measurement of presented blots. The mean ratios of total ubiquitin conjugates to GAPDH ± SEM is presented. n = 3. *, P < 0.05 *vs.* shLacz; #, P < 0.05 *vs.* shCapn1; §, P < 0.05 *vs.* shAtad1. (E) Cleavage of desmin filaments by calpain-1 is facilitated in the presence of ATAD1. The insoluble fraction from 3 d denervated muscle was subjected to cleavage by recombinant calpain-1 in the presence (lanes 5-8) or absence (lanes 1-4) of ATAD1 (ATAD1 was immunoprecipitated from mouse muscle). (F) *Atad1* downregulation promotes normal muscle growth. Measurements of cross-sectional areas of 423 fibers expressing shAtad1 (green bars) versus 423 non-transfected fibers (black bars) in the same muscle. n = 3 mice. Bar: 50 µm. (G) (G)Downregulation of *Atad1* in normal muscle causes accumulation of ubiquitinated desmin filaments. Left: insoluble fractions from normal muscles expressing shLacz or shAtad1 were analyzed by immunoblotting. Right: densitometric measurement of presented blots. Mean ± SEM. n = 4. *, P < 0.05 *vs.* shLacz. (H) Downregulation of *ATAD1* in normal muscle causes accumulation of ubiquitinated soluble proteins. Left: homogenates from normal muscles expressing shLacz or shAtad1 were analyzed by immunoblotting. Right: densitometric measurement of presented blots. The mean ratios of total ubiquitin conjugates to GAPDH ± SEM is presented. n = 4. *, P < 0.05 *vs.* shLacz.

To test this idea more directly, we assayed *in vitro* the susceptibility of phosphorylated desmin filaments to cleavage by calpain-1 in the presence of ATAD1. The insoluble fraction from 3 d denervated muscles, in which GSK3-β was catalyzing desmin IF phosphorylation, was isolated and subjected to cleavage by recombinant calpain-1, with or without the addition of ATAD1. ATAD1 was immunoprecipitated from 7 d denervated muscles, when it is found bound to calpain-1 and other critical interacting partners (Fig. 3C-E). As we had shown before (Aweida et al., 2018), desmin filaments were efficiently processed by calpain-1 *in vitro* as indicated by the appearance of short desmin fragments over time (Fig. 4E, lanes 2-4). However, in the presence of ATAD1, cleavage of phosphorylated desmin filaments by calpain-1 was substantially more efficient, as even smaller fragments of desmin appeared in shorter incubation times, which correlated with a rapid reduction in the amount of phosphorylated desmin (Fig. 4E, compare lanes 6-7 and 2-3). In addition, by 10 min of incubation, phosphorylated monomeric desmin and its cleavage products were virtually absent in the reaction containing ATAD1, most likely due to their processive cleavage by calpain-1 (Fig. 4E, compare lanes 8 and 4), suggesting that desmin is more sensitive to cleavage by calpain-1 in the presence of ATAD1. Thus, on denervation, desmin filament depolymerization by calpain-1 is facilitated by ATAD1.

### *Atad1* downregulation attenuates basal turnover of desmin filaments and of soluble proteins

Because downregulation of *Atad1* reduced atrophy on denervation, we determined whether it also affects the basal turnover of muscle proteins. Downregulation of *ATAD1* by *in vivo* electroporation of shAtad1 into normal TA muscles induced muscle fiber growth because the cross-sectional area of 423 transfected fibers (also express GFP) was significantly larger than that of the surrounding 423 non-transfected ones (Fig. 4F and Table I). ATAD1 attenuated normal muscle growth most likely by promoting the loss of desmin filaments and of soluble proteins because in normal muscles expressing shAtad1 phosphorylated desmin filaments accumulated as ubiquitinated species (Fig. 4G) and ubiquitinated proteins also accumulated in the cytosol (Fig. 4H). Thus, ATAD1 seems to function in normal postnatal muscle to limit fiber growth, and suppression of its activity alone can induce muscle hypertrophy.

## DISCUSSION

In this study we uncovered a new critical cellular role for ATAD1, a AAA-ATPase with previously unknown cofactors and very few identified roles. We identified and validated *in vivo* two interacting partners for ATAD1, UBXN4 and PLAA, and revealed that the ATAD1-PLAA-UBXN4 complex regulates muscle mass by promoting desmin filaments depolymerization and loss. On denervation, ATAD1 binds and cooperates with calpain-1 to facilitate desmin IF depolymerization, and both enzymes are essential for desmin loss. These enzymes are also necessary for the degradation of ubiquitinated proteins in the cytosol of denervated muscles, or their actions allow other proteolytic enzymes to act. In either case, ATAD1 and calpain-1 are clearly general enzymes affecting insoluble and soluble proteins in an additive manner. Moreover, our studies implicate ATAD1 in the slower turnover of desmin IF and soluble proteins in normal muscle. Consequently, suppression of ATAD1 activity alone not only attenuates atrophy but can also induce muscle growth.

Reducing ATAD1 or calpain-1 (Aweida et al., 2018) functions by shRNA attenuated the decrease in fiber diameter on denervation. The stabilization of desmin IF in these muscles coincided with accumulation of ubiquitinated proteins in the cytosol, consistent with our prior findings demonstrating that desmin IF loss in atrophy promotes overall proteolysis (Aweida et al., 2018; Volodin et al., 2017; Cohen et al., 2012). The reduced structural integrity of desmin filaments on denervation is likely the key step in the destabilization of insoluble proteins (e.g. myofibrils) during atrophy, leading to the enhanced solubilization and degradation in the cytosol. Degradation of soluble desmin in the cytosol also requires ATAD1 (Fig. 2C). Such soluble pools of desmin and its homolog vimentin are small as these proteins mostly exist in the cell assembled within filaments (Soellner et al., 1985; Cohen et al., 2012; Aweida et al., 2018). Possibly, this soluble pool of desmin functions either as precursors to the mature filament or as components released during filament turnover. Because desmin IF disassembly is prevented in atrophying muscles expressing shAtad1, the soluble desmin that accumulates in the cytosol of these muscles likely represents new precursors to the mature filament. In addition to the essential roles of ATAD1 and calpain-1 in atrophy, the accelerated proteolysis during atrophy seems to require both enzymes and additional cofactors (e.g. UBXN4, PLAA) to function with them, as well as posttranslational modifications to enhance the susceptibility of desmin filaments to these enzymes. For example, desmin filaments must be modified by phosphorylation before their disassembly by ATAD1 and calpain-1 (Fig. 2F) (Aweida et al., 2018).

Because ATAD1 is important for muscle atrophy and for normal muscle growth, identification of its cofactors was essential to understand the mechanisms of protein homeostasis and loss. We used three independent mass spectrometry-based proteomics approaches and identified a number of UPS components as novel interacting partners for ATAD1 *in vivo*, all of which are important for protein degradation by the proteasome (Table II). Using these analyses, we identified and validated UBXN4 and PLAA as ATAD1 partners, and the ATAD1-PLAA-UBXN4 complex as responsible for depolymerization and loss of ubiquitinated desmin IF in skeletal muscle. Precisely how ubiquitination enables ATAD1-driven desmin IF disassembly is an important question for further research. Although ATAD1 can bind ubiquitin (Fig. 3C and Table II), we found that this AAA-ATPase associates with phosphorylated desmin, whether they are polyubiquitinated (e.g., in atrophying muscles) or not (e.g., in atrophying muscles expressing shTrim32), suggesting that ATAD1 binds substrates either before or during their polyubiquitination. In fact, ATAD1 accumulates on desmin filaments in muscles expressing shTrim32 suggesting that ubiquitination by TRIM32 is not required for ATAD1 binding but is rather important for its release from desmin IF. In other words, ubiquitination by TRIM32 is likely important for ATAD1-mediated extraction of ubiquitinated desmin from the insoluble filaments into the cytosol, where desmin is degraded by the proteasome (Fig. 2D). TRIM32 was not identified as ATAD1 partner by our mass spectrometry analyses, suggesting that the association of this E3 with desmin filaments is not dependent on ATAD1. It will be important to determine if ubiquitination by TRIM32 is sufficient for ATAD1-mediated disassembly of desmin filaments, or whether the ubiquitin chains generated by TRIM32 are extended further by ubiquitin ligases that are recruited by ATAD1 to desmin (e.g. HUWE1, see Table II). Desmin was recently reported to be ubiquitinated by the ubiquitin ligase MURF1 in normal mouse muscles overexpressing *Murf1*, though this ubiquitination of desmin did not lead to degradation (Baehr et al., 2021). We could not precipitate MURF1 with ATAD1 from denervated muscle extracts, and it remains to be determined if MURF1 acts on desmin *in vivo* during atrophy.

It is noteworthy that co-fractionation with ATAD1 using three approaches did not identify the exact same subset of putative adaptors (Table II), suggesting a dynamic association of ATAD1 with its cofactors. It is also possible that distinct ATAD1’s interacting partners may have been missed by our analyses, especially if specific stimuli are required to facilitate such interaction. Future studies will determine how these adaptor proteins function with ATAD1 to promote substrate recruitment and processing, and whether their function is regulated by substrates or other cofactors. These various binding partners could be precipitated with ATAD1 from the cytosolic fraction, and could also bind desmin in the insoluble pellet of normal and atrophying muscles (Table II). ATAD1 appears to be recruited to desmin filaments early during atrophy to promote desmin release during filament turnover. This AAA-ATPase complex clearly binds desmin at 3 d after denervation when desmin filaments are intact, and 4 days later, when desmin IF solubilization is rapid (Volodin et al., 2017; Aweida et al., 2018), ATAD1 and its binding partners accumulate in the cytosol, bound to soluble ubiquitinated desmin (Fig. 3D). In addition, ATAD1 catalyzes desmin IF disassembly only when these filaments are phosphorylated, presenting ATAD1 as the only known AAA-ATPase that acts preferentially on phosphorylated substrates. It is likely that ATAD1 interacting partners, which contain a phospho-Serine/Threonine binding domain, mediate ATAD1 association with phosphorylated desmin IF. Although calpain-1 binds preferentially to phosphorylated desmin IF (Aweida et al., 2018) and is present in ATAD1 complex, it is not required for ATAD1 recruitment to desmin IF because in denervated muscles expressing shCapn1, ATAD1 accumulated on desmin filaments (Fig. S1). Thus, calpain-1 facilitates the release of ATAD1 from desmin IF, which is consistent with the cooperative roles of these enzymes in promoting desmin filament disassembly.

The previously known cellular roles for ATAD1 are regulation of synaptic plasticity in the brain (Zhang et al., 2011; Li et al., 2016), and extraction of mislocalized proteins from mitochondrial membrane (Li et al., 2019; Wohlever et al., 2017). Our findings that ATAD1 disassembles ubiquitinated intermediate filaments in skeletal muscle present a novel function for this AAA-ATPase complex, which is similar to p97/VCP complex that also extracts ubiquitinated proteins from larger structures (Stach and Freemont, 2017; Volodin et al., 2017; Piccirillo and Goldberg, 2012). ATAD1 is expressed in various tissues (Fig. S2) and may thus have many cellular roles that are probably dictated by the specific cofactors that it binds. This study is the first to identify ATAD1’s binding partners, and to demonstrate that ATAD1-PLAA-UBXN4 complex binds the insoluble phosphorylated desmin IF and promote the release of ubiquitinated species into the cytosol. Accordingly, downregulation of either *ATAD1, PLAA* or *UBXN4* was sufficient to prevent solubilization and loss of ubiquitinated desmin filaments. PLAA and UBXN4 are also known cofactors for p97/VCP (Liang et al., 2006; Papadopoulos et al., 2017), a AAA-ATPase that was not in our datasets, indicating that p97/VCP adaptors can bind and function with other AAA-ATPases. It is plausible that ATAD1 binds additional factors, as our mass spectrometry data strongly suggest, whose specific cellular roles merit further study. It is also likely that such factors will be required for ATAD1-mediated removal of mislocalized proteins from the mitochondrial membrane (Wohlever et al., 2017). For example, UBXN4 contains a putative transmembrane segment (Sasagawa et al., 2010), and via association with ATAD1 may anchor and recruit ATAD1 complex to mitochondrial membrane proteins. Mitochondria are located in close proximity to the myofibrils in muscle, and are linked to each other and to other cellular organelles via desmin IF (Milner et al., 2000). Thus, in skeletal muscle the ATAD1-PLAA-UBXN4 complex may serve a dual role in the extraction of proteins from desmin filaments and the mitochondria.

The strongest evidence for the importance of ATAD1 and calpain-1 cooperation for the accelerated loss of desmin was our finding that simultaneous downregulation of both enzymes inhibited desmin loss to a greater extent than when each enzyme was downregulated alone. In addition, our data indicate that desmin filaments are more sensitive to cleavage by calpain-1 *in vitro* in the presence of ATAD1 than in samples containing calpain-1 alone. Thus, the present findings uncover a novel mechanism involving a cooperation between an AAA-ATPase and a protease to facilitate filament loss. Because ATAD1 and calpain-1 also affect ubiquitinated soluble proteins in an additive fashion (Fig. 4D), and since these enzymes are ubiquitously expressed, this new mechanism probably contributes to proteolysis in other cells, and may also apply to other AAA-ATPases and proteases.

Desmin loss is a critical event leading to myofibril destruction and muscle atrophy, therefore, identifying key regulators of this process should be of major therapeutic promise to treat muscle wasting. ATAD1, along with the previously identified key enzymes the protein kinase GSK3-β, the ubiquitin ligase TRIM32, and the Ca^2+^-specific protease calpain-1, appear to act on the desmin cytoskeleton in a specific order to promote its loss (Aweida et al., 2018). We propose that ATAD1 binds desmin filaments early after denervation (at 3 d), and together with its interacting partners promote a slow dissociation of desmin filaments (Fig. 5). Then, at 7 d after denervation, when Ca^2+^ levels rise and calpain-1 is activated, desmin depolymerization is accelerated by the cooperative functions of calpain-1 and ATAD1. We propose that these two enzymes function in a “Pull and Chop” mechanism, in which the ATAD1 complex extracts ubiquitinated desmin filaments, exposing on desmin cleavage sites for calpain-1, consequently facilitating desmin cleavage, solubilization and degradation by the proteasome. Cleavage by calpain-1 may expose degrons on desmin to facilitate its degradation, as has been proposed for other proteases (Varshavsky, 2019). Such cooperation may be important to facilitate protein degradation during rapid types of atrophy, for example during fasting or denervation, where muscle loss occurs over days, and may be less relevant in prolonged types of atrophy such as during aging where muscle loss is slow and gradual.

**Figure 5:**
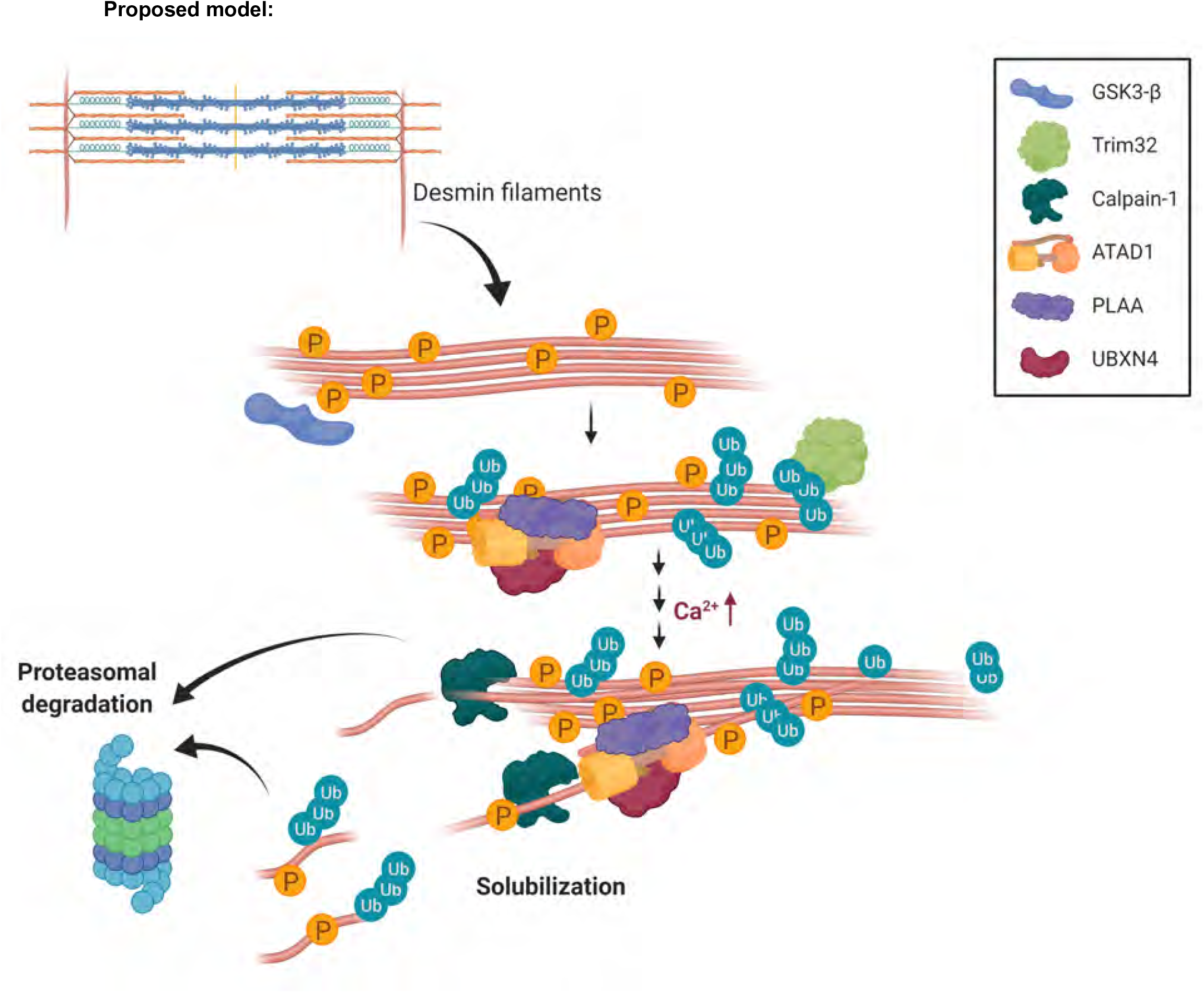
Proposed mechanism for desmin filament loss during denervation-induced atrophy. Early after denervation (at 3 d), ATAD1 binds phosphorylated and ubiquitinated desmin filaments and together with its interacting partners promote a slow dissociation of desmin IF. Then, at a more delayed phase, when calcium levels rise and calpain-1 is activated, desmin filament depolymerization is accelerated by the cooperative functions of calpain-1 and ATAD1. We propose that these two enzymes function in a “Pull and Chop” mechanism, in which the ATAD1 complex extracts ubiquitinated desmin filaments, exposing on desmin cleavage sites for calpain-1, consequently facilitating desmin cleavage, solubilization and subsequent degradation by the proteasome. The illustration was created with BioRender.com">BioRender.com.

## MATERIALS AND METHODS

### Animal work

All animal experiments were consistent with Israel Council on Animal Experiments guidelines and the Institutional regulations of Animal Care and Use. Specialized personnel provided mice care in the institutional animal facility. We used adult CD-1 male mice (∼30 g) (Envigo) in all experiments. Muscle denervation was performed by sectioning the sciatic nerve on one limb, while the contralateral leg served a control. Muscles were excised at 3, 7, 10 or 14 days after denervation. To inhibit proteasome activity, mice were injected with Bortezomib (3mg/kg body weight, i.p.) or DMSO, and an hour later were sacrificed.

### Antibodies, constructs and materials

Oligos encoding shRNA against *ATAD1, PLAA, and UBXN4* were cloned into pcDNA 6.2GW/EmGFP-miR vector using Invitrogen’s BLOCK-iT RNAi expression kit as described (Goldbraikh et al., 2020; Volodin et al., 2017). shCapn1, shTrim32, shLacz and the plasmid encoding HA-tagged GSK3-β dominant negative (K85A catalytically dead mutant) were described and validated before (Aweida et al., 2018). A plasmid encoding 6His-Calpain-1 was a kind gift from Dr. Shoji Hata, Tokyo Metropolitan Institute of Medical Science, Tokyo, JAPAN. ATAD1 (for immunofluorescence and immunoblotting) and anti PLAA (for immunofluorescence) antibodies were from Abcam. Desmin antibodies were from Abcam and Developmental Studies Hybridoma bank (developed by D.A. Fischman, Cornell University Medical College). GAPDH, UBXN4, ATAD1 (for immunofluorescence), calpain-1 (for immunofluorescence) and laminin antibodies were from Sigma, calpain-1 (for immunoblotting) from Cell Signaling, anti-phospho-Threonine/Serine from ECM Biosciences, and anti-PLAA from Invitrogen. Anti-mono and polyubiquitin conjugates was from Enzo (#BML-PW1210). TRIM32 antibody was kindly provided by Dr. Knoblich (Institute of Molecular Biotechnology, Vienna, Austria).

### *In vivo* transfection

*In vivo* electroporation experiments were performed in adult CD-1 male mice (∼30 g) as reported (Goldbraikh et al., 2020; JE et al., 2021). Briefly, 20 µg of plasmid DNA was injected into adult mouse TA muscles, and a mild electric pulse was applied using two electrodes (12 V, 5 pulses, 200 ms intervals). For fiber size analysis, muscle cross-sections were fixed in 4 % paraformaldehyde (PFA) and fiber membrane was stained with laminin antibody (1:50, see details under immunofluorescence). Cross sectional areas of muscle sections (20 μm) were analyzed using Imaris software (Bitplane). Images were collected using a Nikon Ni-U upright fluorescence microscope with Plan Fluor 20 x 0.5 NA objective lens and a Hamamatsu C8484-03 cooled CCD camera, at room temperature. For biochemical analyses, muscles that are at least 60-70% transfected were used.

### Fractionation of muscle tissue

To obtain whole cell extracts, muscles were homogenized in lysis buffer (20 mM Tris pH 7.6, 5 mM EGTA, 100 mM KCl, 1% Triton X-100, 1 mM PMSF, 10 mM Sodium Pyrophosphate, 3 mM Benzamidine, 10 μg/ml Leupeptin, 10 μg/ml Aprotinin, 50 mM NaF and 2 mM Sodium OrthoVanadate), and incubated for 1 hr at 4°C. After centrifugation at 6000 x *g* for 20 mins at 4° C, the supernatant (i.e. soluble fraction) was stored at −80° C. The insoluble pellet was washed once with homogenization buffer and twice with suspension buffer (20 mM Tris pH 7.6, 100 mM KCl, 1 mM DTT and 1 mM PMSF), and after a final centrifugation at 6000 x *g* for 10 mins at 4°C, the insoluble pellet (i.e. purified myofibrils and desmin filaments) was re-suspended in storage buffer (20 mM Tris pH 7.6, 100 mM KCl, 1 mM DTT and 20 % glycerol) and kept at −80°C. To isolate desmin filaments, 30μg of the insoluble muscle pellet was resuspended in ice-cold extraction buffer (0.6 M KCl, 1% Triton X-100, 2 mM EDTA, 2 mM PMSF, 1 x PBS, 10 μg/ml leupeptin, 3 mM benzamidine, 10 μg/ml Aprotinin, 50 mM NaF, 2 mM Sodium OrthoVanadate and 10 mM Sodium Pyrophosphate) for 10 mins on ice. After centrifugation at 6000 x *g* for 10 mins at 4° C, the pellet was resuspended in 20 mM Tris pH 7.6 and analyzed by SDS-PAGE and immunoblotting.

### Protein analysis

For immunoblotting, soluble (25 μg) or insoluble (3 μg) fractions were resolved by SDS-PAGE, transferred onto PVDF membranes and immunoblotted with specific primary antibodies, and secondary antibodies conjugated to HRP. For immunoprecipitation, 300 μg of muscle soluble fraction were incubated with a specific primary antibody (the control sample contained non-specific IgG) overnight at 4 °C, and protein A/G agarose was then added for 4 hr. To remove nonspecific or weakly associated proteins, the precipitates were washed extensively with 10 bed volumes of each of the following buffers: buffer A (50 mM Tris-HCl, pH 8, 500 mM NaCl, 0.1% SDS, 0.1% Triton, 5 mM EDTA), buffer B (50 mM Tris-HCl, pH 8, 150 mM NaCl, 0.1% SDS, 0.1% Triton, 5 mM EDTA) and buffer C (50 mM Tris-HCl, pH 8, 0.1% Triton, 5 mM EDTA). Protein precipitates were eluted with protein loading buffer containing 50 mM DTT, and were analyzed by immunoblotting.

### Immunofluorescence labeling of frozen muscle sections

Frozen longitudinal sections of mouse TA were cut at 10 μm, fixed in 4% PFA for 10 min at room temperature (RT), and washed in PBST (PBS, 0.1% Triton X-100) three times. For antigen retrieval, slides were incubated in 10mM Citrate buffer (9ml of 0.1M Citric acid, 41ml of 0.1M Sodium citrate, 450ml water, pH 6.0) on low heat (lowest power in microwave) for 20 min, and then left to cool down at RT (∼ 2 hr). Following one wash in PBST, the slides were incubated in blocking solution (0.2% BSA, 5% goat serum and 5% donkey serum in PBST) for 1 hr at RT. Immunofluorescence was performed using ATAD1 (1:50), Calpain-1 (1:50), PLAA (1:20) or UBXN4 (1:50) antibodies overnight at 4 °C, followed by 2 hr incubation at RT with secondary antibodies conjugated to Alexa Fluor 568 or 647 (1:400). Primary antibodies were diluted in blocking solution, and secondary antibodies were diluted in 0.2% BSA/PBST. Sections were imaged using the Elyra 7 eLS lattice SIM super resolution microscope by Zeiss with a pco.edge sCMOS camera. A X63 1.46 NA oil immersion objective with 561nm and 642nm lasers. 16-bit 2D image data sets were collected with 13 phases. The SIM^2 image processing tool by Zeiss was used.

### Colocalization analysis by Imaris

Colocalization measurements were performed using the Imaris software (Bitplane, ver 9.31), after setting an appropriate threshold that was kept constant during the entire analysis. At least three images per protein staining were analyzed. Imaris segmented the stained proteins with the “Spots” module. The spots were set to 100 nm in diameter that is below the light diffraction limit but above the SIM^2 super resolution imaging limit (60 nm). Finally, colocalized spots were identified by using imaris extension “coloc spots” and a distance threshold between close spots was set to 100 nm from each other.

### Purification of recombinant calpain-1

6His-tagged calpain-1 was purified from BL21 bacteria (OD 0.6) grown with 0.2 mM IPTG at 17°C overnight. After centrifugation (4000 *g* at 4°C for 20 min), the pellet was lysed with lysis buffer (50 mM NaH2PO4, 300 mM NaCl, and 10 mM imidazole) and a microfluidizer, and centrifuged at 10,000 *g* at 4°C for 20 min. The obtained supernatant was incubated with a nickel column for 1 hr at 4°C, the column was washed twice with wash buffer (50 mM NaH2PO4, 300 mM NaCl, and 20 mM imidazole), and 6His-calpain-1 was eluted with elution buffer (50 mM NaH2PO4, 300 mM NaCl, and 250 mM imidazole). After dialysis against 50 mM Tris, pH 7.6, overnight at 4°C, 1 mM DTT and 10% glycerol were added, and purified calpain-1 was stored at −80°C.

### *In vitro* cleavage assay by calpain-1

ATAD1 was immunoprecipitated from 300 μg of soluble fraction from 7 d denervated TA muscle (see Protein analysis). Two sets of tubes containing 30 μg of the myofibrillar fraction (see Fractionation of muscle tissue) from denervated muscles (3 d), were incubated with 6.5 μg purified 6His-Calpain-1 in reaction buffer (100 mM Hepes, 10 mM DTT and 5 mM CaCl_2_) for 2, 5 or 10 mins at 30° C, in the presence or absence of ATAD1. Negative control samples also contained 50 mM EGTA. Desmin cleavage by calpain-1 was assessed by SDS-PAGE and immunoblotting.

### Identification of ATAD1 interacting partners using SEC

Seven days denervated lower limb muscles (1 g) were homogenized in 10 volumes of 1 % PFA/PBS (v/w), and incubated for 10 min at RT. The reaction was quenched by the addition of 0.125 M glycine for 5 min at RT, centrifuged at 3000 x *g* for 5 min at 4°C, and the supernatant was adjusted to 1X homogenization buffer (see fractionation of muscle tissue. Adjustment was performed using 10X homogenization buffer). Then, proteins in the supernatant were precipitated with 40 % ammonium sulfate and centrifuged at 20,000 x *g* for 30 min at 4° C. The pellet was resuspended in 0.5 ml of buffer A (50 mM Tris pH 8, 0.5 M NaCl) 5 mM EGTA, 1% Triton X-100, 1 mM PMSF, 10 mM Sodium Pyrophosphate, 3 mM Benzamidine, 10 μg/ml Leupeptin, 10 μg/ml Aprotinin, 50 mM NaF and 2 mM Sodium OrthoVanadate), and loaded onto a Superdex 200 10/300 Gel Filtration column (GE Healthcare Life Sciences, UK) equilibrated with 50 mM Tris pH 8, 0.5 M NaCl. Proteins were eluted at a flow rate of 0.5 ml/min and 500 μl fractions were collected. Even fractions were analyzed by silver staining and immunoblotting.

### Protein identification by mass spectrometry

For mass spectrometry analysis, protein bands were excised from gel, reduced with 3 mM DTT in 100 mM ammonium bicarbonate [ABC] (60°C, 30 mins), modified with 10mM Iodoacetamide in 100 mM ABC (at the dark, RT, 30 mins) and digested in 10% Acetonitrile and 10 mM ABC with modified trypsin (Promega) at a 1:10 enzyme-to-substrate ratio overnight at 37°C. The resulted peptides were desalted using C18 tips (Homemade stage tips, Empore), dried, and re-suspended in 0.1% Formic acid. The peptides were then resolved by reverse-phase chromatography on 0.075 × 180-mm fused silica capillaries (J&W) packed with Reprosil reversed phase material (Dr Maisch GmbH, Germany), and were eluted with linear 60 mins gradient of 5 to 28 % 15 mins gradient of 28 to 95 % and 15 mins at 95 % acetonitrile with 0.1 % formic acid in water at flow rates of 0.15 μl/min. Mass spectrometry was performed by Q Exactive plus mass spectrometer (Thermo) in a positive mode using repetitively full MS scan followed by collision induces dissociation (HCD) of the 10 most dominant ions selected from the first MS scan.

The mass spectrometry data was analyzed using Proteome Discoverer 1.4 software with Sequest (Thermo) and Mascot (Matrix Science) algorithms against mouse uniport database with mass tolerance of 10 ppm for the precursor masses and 0.05 amu for the fragment ions. Minimal peptide length was set to six amino acids and a maximum of two mis-cleavages was allowed. Peptide-and protein-level false discovery rates (FDRs) were filtered to 1 % using the target-decoy strategy. Protein tables were filtered to eliminate the identifications from the reverse database and from common contaminants. Semi quantitation was done by calculating the peak area of each peptide based its extracted ion currents (XICs) and the area of the protein is the average of the three most intense peptides from each protein.

### Real-Time qPCR

Total RNA was isolated from muscle using TRI reagent (T9424; Sigma-Aldrich) and served as a template for the synthesis of cDNA by reverse transcription. Real-time qPCR was performed on mouse target genes using specific primers (Table S1) and the Perfecta SYBR Green qPCR Kit (95073-012; Quanta Biosciences) according to the manufacturer’s protocol.

### Statistical analysis and image acquisition

Data are presented as mean ± SEM. The statistical significance was accessed with one-tailed Student’s *t* test. Muscle sections for fiber size analysis were imaged at room temperature with a Nikon Ni-U upright fluorescence microscope with Plan Fluor 20x 0.5NA objective lens and a Hamamatsu C8484-03 cooled CCD camera. Image acquisition and processing was performed using the Metamorph or the Imaris software (Bitplane). Statistics on fiber size distributions was performed using A-statistics and Brunner-Manzel test as described in our recent methodology paper (JE et al., 2021). Muscle sections in Fig. 3E were imaged using the Elyra 7 eLS lattice SIM super resolution microscope by Zeiss with a pco.edge sCMOS camera. A X63 1.46 NA oil immersion objective with 561nm and 642nm lasers. 16-bit 2D image data sets were collected with 13 phases. The SIM^2 image processing tool by Zeiss was used. Black and white images were processed with Adobe Photoshop CS5, version 12.1×64 software. Quantity One algorithm (Bio-Rad Laboratories, version 29.0) was used for densitometric measurements of intensity of protein bands.

## ACKNOWLEDGEMENTS

This project was supported by grants from the Israel Science Foundation (grant no. 1068/19), Israel Ministry of Health (grant no. 3-0000-16061), and the Niedersachsen-Deutsche (grant no. ZN3008) to S. Cohen. Additional funds were received from the Russell Berrie Nanotechnology Institute, Technion to S. Cohen.

We thank the Smoler Proteomics Center at Technion for the mass spectrometry analysis. The illustration in Fig. 5 was created with BioRender.com.

The authors declare no competing financial interests.

## AUTHOR CONTRIBUTIONS

D.A. performed all experiments. D.A., S.C designed experiments, analyzed data, and wrote the paper.

## SUPPLEMENTARY MATERIAL

**Figure S1:**
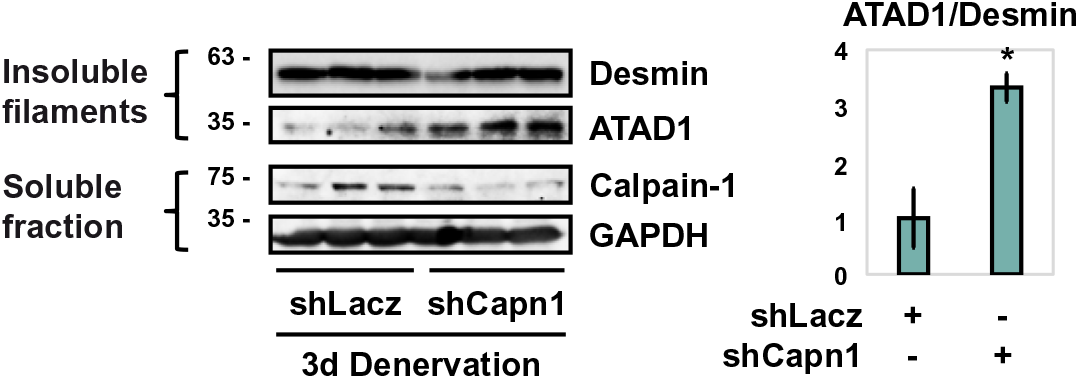
Calpain-1 is not required for ATAD1 recruitment to desmin filaments, but rather to ATAD1 release when desmin IF depolymerize. Left: desmin filaments isolated from denervated muscles expressing shLacz or shCapn1 were analyzed by immunoblotting. Right: densitometric measurement of presented blots. Mean ratio of ATAD1 to desmin ± SEM is presented. n = 3. *, P < 0.05 *vs.* shLacz.

**Figure S2:**
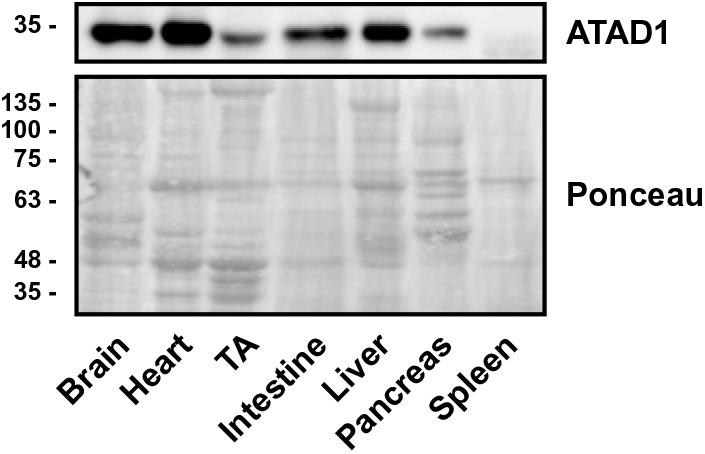
ATAD1 is expressed in various tissues. Soluble fractions from brain, heart, TA muscle, intestine, liver, pancreas and spleen were analyzed by SDS-PAGE and immunoblotting.

**Table S1.**
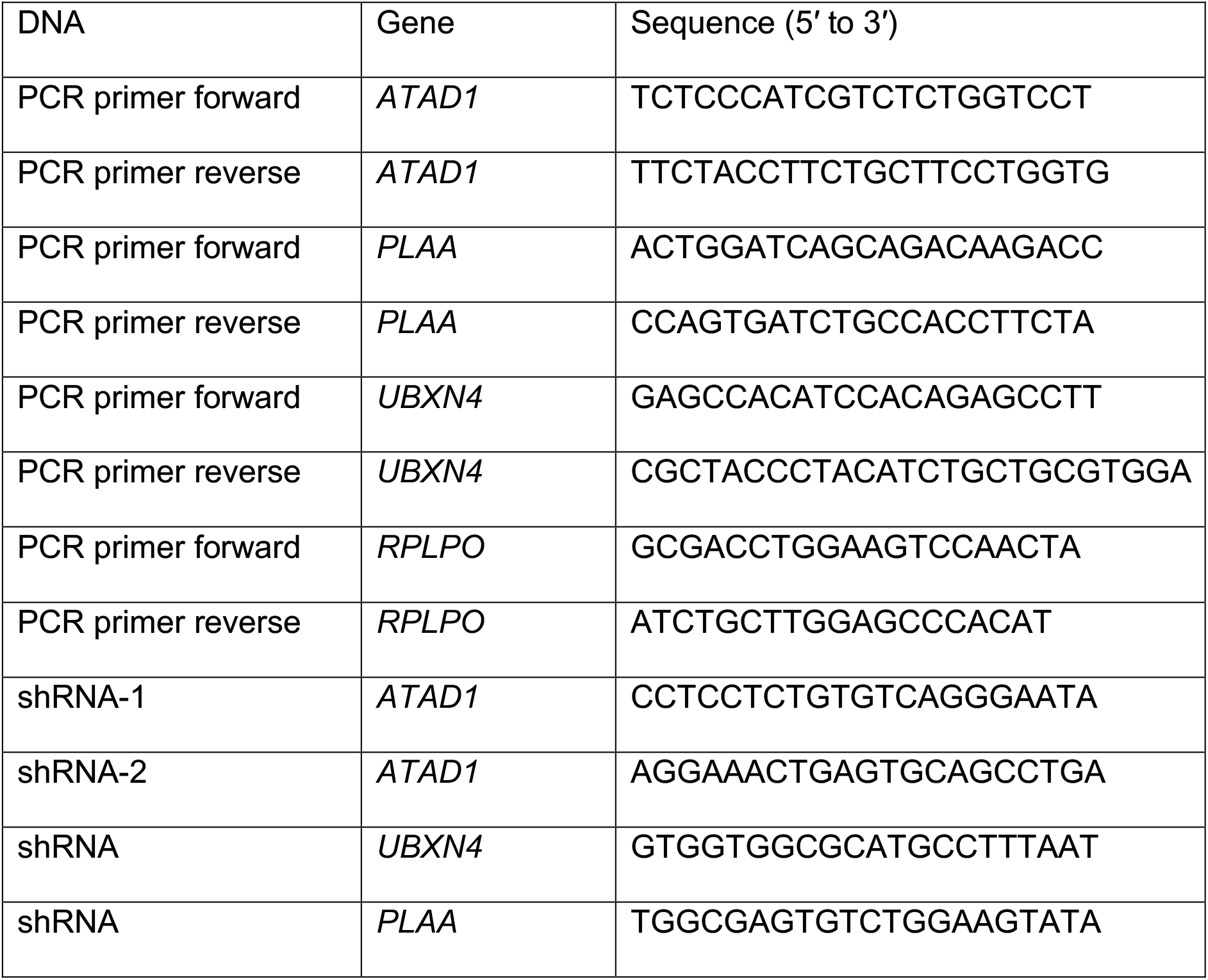
Quantitative PCR primers and shRNA oligonucleotides used in the present study.

